# Fine oculomotor knowledge enhances high-acuity vision

**DOI:** 10.1101/2022.03.17.483854

**Authors:** Zhetuo Zhao, Ehud Ahissar, Jonathan D. Victor, Michele Rucci

## Abstract

It has long been debated how humans resolve fine details and perceive a stable visual world despite the fixational motion of their eyes, the incessant ocular jitter that occurs in the intervals between voluntary gaze shifts. Current theories assume these processes to rely solely on the visual input to the retina, without contributions from motor and/or proprioceptive sources. Here we show that contrary to this widespread assumption, the visual system has access to high-resolution extra-retinal knowledge of fixational eye motion and uses it to deduce spatial relations. Building on recent advances in gaze-contingent display control, we created a spatial discrimination task in which the stimulus configuration was entirely determined by oculomotor activity. Our results show that humans correctly infer geometrical relations even when no spatial information is delivered to the retina and accurately combine high-resolution extraretinal monitoring of gaze displacement with retinal signals. These findings reveal a multimodal strategy for encoding spatial details, in which fine oculomotor knowledge is used to interpret the fixational input to the retina.

## Introduction

Our eyes are never at rest. Since fine visual resolution is restricted to a tiny portion of the retina, the fovea, humans use eye movements to inspect objects of interest. Remarkably, the eyes remain in motion even in the intervals between voluntary gaze shifts, the so-called “fixation” periods in which visual information is acquired and processed. In these periods, a persistent eye jitter, known as ocular drift, continually perturbs the direction of gaze, moving the projection of the stimulus on the retina across dozens of receptors.

Given the extent of ocular drift and the temporal responses of retinal neurons, it has long been questioned how the visual system manages to avoid perceptual blurring during fixation and establish stable high-acuity representations^1–4^. Multiple theories have been proposed. Some regard the fixational motion of the eye as a challenge to be overcome through specific decoding strategies^5,6^. Others argue that eye movements are beneficial for processing spatial information, either by transforming spatial patterns into temporal modulations^7–10^ or by following spatial registration strategies similar to those used in computer vision to enhance image resolution^11,12^. Although the proposed theories differ widely in their specific mechanisms, they all share the common assumption that spatial representations at fixation are established solely based on the visual input signals impinging onto the retina, without making use of information from other sources.

This standard assumption, however, contrasts with the multimodal and sensorimotor integration that is known to occur in the presence of larger eye movements, such as the rapid gaze shifts (saccades) and tracking movements (smooth pursuits) that bring and maintain objects onto the fovea. With these movements, interpretation of retinal activity critically depends on motor and proprioceptive knowledge about how the eyes move. Extraretinal signals are known to modulate visual responses by both enhancing and attenuating sensitivity, often in a dynamic manner at specific times during the movements^13–22^. Extraretinal modulations are deemed to be essential for extracting information from the retinal flow^23,24^, establishing spatial representations^25–29^, and discarding the motion of the retinal image caused by the eye movements themselves^30,31^.

Various factors have contributed to the current tenet that a similar visuomotor integration does not take place during fixation. From a historical perspective, vision science has traditionally approximated the fixational input to the retina as an image, neglecting the incessant motion of the eye and/or assuming this motion to be too small to yield reliable motor or proprioceptive signal. The eyes appear to wander erratically during fixation, leading many researchers to conclude that ocular drift stems from limits in oculomotor control ^32,33^ and is, thus, unlikely to be monitored. Reinforcing this idea, previous attempts to identify extraretinal signals associated with fixational drifts reported negative results^34,35^, and several studies have argued that retinal signals are solely responsible for establishing stable visual representations during fixation (*e*.*g*., [36]).

However, contrary to the mainstream assumption, it has long been proposed that ocular drift may actually represent a form of slow control aimed at delivering a desired amount of retinal image motion^37,38^. This proposal has received renewed support from recent findings, including the observation that drift partly counteracts the physiological instability of the head^39^, as well as task-dependent changes in drift characteristics^40–42^. Furthermore, previous studies that searched for motor knowledge of fixational drifts either did not control for the spatial information delivered to the retina^34^ or focused on relatively long temporal windows, intervals over which memory decays could have played a role (*e*.*g*., [35]). These considerations raise the need for more specific investigations on the mechanisms by which stable high-resolution spatial representations are established during the incessant fixational motion of the eye.

Here we built upon recent advances on high-resolution eye-tracking and gaze-contingent display control, the capability to modify the stimulus in real-time according to the observer’s eye movements, to precisely control retinal stimulation. We developed a spatial discrimination task that cannot be accomplished solely based on the visual input signals to the retina, but rather, depended critically on knowledge of eye position. We show that despite the lack of spatial information in the retinal input, the visual system is capable of reconstructing the configuration of the stimulus, and therefore estimating the fixational motion of the eye, with exquisite sensitivity. These results show that humans possess fine motor knowledge of the way the eye drifts during fixation and integrate this information into high-resolution spatial representations.

## Results

We developed a task that requires motor knowledge of the direction in which the eye moves to be successfully executed. In a forced-choice task, subjects discriminated the spatial configuration of a stimulus that entirely depended on their performed eye movements. They reported whether the bottom bar of a Vernier appeared to be to the right or left of the top bar (Fig. 1A), but, unlike a conventional spatial judgment, the two bars of the Vernier were never visible simultaneously, and no information about their spatial offset was ever delivered to the retina. This was achieved via a gaze-contingent procedure that rendered the stimulus on the display as seen through a retinally-stabilized aperture, a thin slit that moved under real-time computer control together with the eye to restrict stimulation to a narrow vertical strip on the retina centered on the fovea (Fig. 1B). In this way, as the normal fixational motion of the eye swept the aperture across the stimulus, the two bars appeared sequentially at vertically aligned positions on the retina (Fig. 1C), yielding input signals that—under ideal conditions—are not informative for the task (Fig. 1D).

**Fig. 1:**
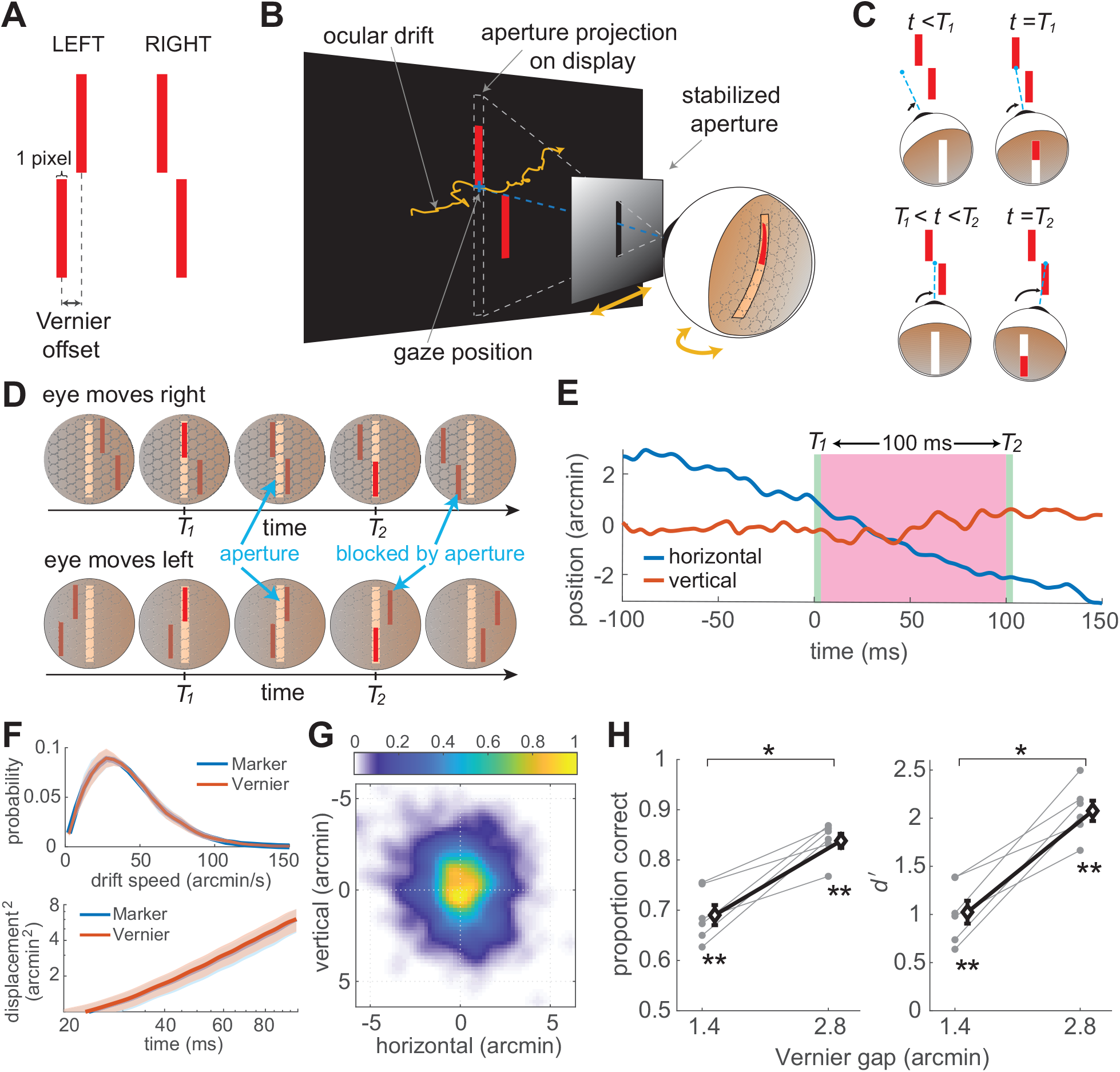
Estimating spatial relations via eye movements. (**A-C**) Experimental design. (*A*) Subjects reported the spatial configuration of a Vernier (left or right) viewed through a retinally-stabilized aperture. (*B*) The aperture moved together with the eye, to allow stimulation of only a thin vertical strip on the retina. The width of the aperture was equal to that of each bar in the Vernier (28′ long; 1.4′, the angle covered by one pixel on the CRT). (*C*) In this way, each Vernier bar was visible only when it directly overlapped with the aperture, resulting in vertically-aligned bar exposures on the retina. (**D**) Motor knowledge of eye movements is required to accomplish this task. The same visual input signals can be obtained with different configurations of the stimulus, when the eye drifts in opposite directions. (**E**) Example trace of eye movements in a trial. The shaded green regions mark the periods of exposure of each Vernier bars. The pink region indicates the inter-stimulus interval (ISI), here 100 ms. (**F-G**) Ocular drift characteristics in the task. (*F*) Mean eye speed and displacement are virtually identical to those measured in the same subjects while fixating on a marker. Shaded regions represent ± one SEM across subjects. (*G*) Average probability distribution of gaze displacement in between bar exposures. (**H**) Subjects correctly reported the configuration of the stimulus. Both proportion correct (left) and d′ (right) were significantly above chance (^*\*\**^*p <* 0.0005, two-tailed t-test) and improved as the Vernier gap increased (^*\**^*p <* 0.0025, paired two-tailed t-test). Bold diamonds and associated error bars represent averages across subjects (*N* = 6) ± one SEM. Gray circles are the individual subjects data.

In practice, unbeknownst to the observer, the two bars were displayed one below and one above the position of the center of gaze at two separate times (*T*_1_ and *T*_2_ in Fig. 1E), the first at a random time from the beginning of the trial, and the second after a fixed delay from the disappearance of the first bar (the inter-stimulus interval, ISI). Subjects moved their eyes normally under these conditions while attempting to maintain fixation at the remembered location of a marker (a 5′ dot) briefly presented at the beginning of each trial. They alternated occasional small saccades with periods of ocular drift, which moved the eye in its stereotypical, seemingly erratic fashion with characteristics similar to those measured from the same observers when maintaining fixation on a visible marker (Fig. 1F). In this study we specifically focused on fixational drifts and discarded all trials in which subjects performed saccades or microsaccades. With an ISI of 100 ms, ocular drift resulted in displacements of the line of sight distributed around ±4′ (Fig. 1G).

Remarkably, subjects were highly proficient in reporting the stimulus configuration, even though its spatial layout was never made explicit on the retina (Fig. 1H). Their qualitative experience consisted of two successive flashes with a clear spatial offset. Performance was significantly above chance already at the smallest Vernier offset that could be presented, a gap of only 1.4′ corresponding to the spacing of just one single pixel on the display. Performance further increased with larger gaps, with a two-fold increment in d′ as the Vernier offset increased to 2.8′. These results were highly consistent across individuals: all subjects were able to successfully accomplish the task. In each individual observer, performance was significantly above chance at all Vernier gaps (*p <*0.021, one-tailed bootstrap test), with the exception of one subject at the smallest offset (1.4′) for which the d′ was close to significance (*p* =0.065).

These results were not caused by possible biases—and thereof knowledge—in the individual direction of eye movements, *i*.*e*., the realization that perhaps drift was more pronounced in one direction. No obvious directional biases were observed in the recorded data, and horizontal displacements in the two directions were approximately symmetrically distributed (Fig. S1A). Furthermore, performance was high in both the trials in which the eye drifted to the left and to the right (Fig. S1B,C), indicating knowledge of the specific direction of ocular drift in each individual trial. Thus, these data suggest that humans have access to high-resolution extra-retinal information of how the eye moves during fixation.

Given these unexpected results, we wondered whether our methods of visual stimulation inadvertently introduced spurious spatial cues. Meticulous care had been taken to eliminate all obvious retinal cues that could inform about the stimulus configuration. This included conducting the experiments in complete darkness—while preventing dark adaptation with brief light exposure between block of trials—to avoid visual references; using a fast-phosphor high-speed CRT display to minimize persistence; and lowering the monitor intensity to minimum settings to ensure that the edges of the monitor were not visible. We questioned, however, whether more subtle cues, such as the baseline luminance of the CRT display or possible residual phosphor persistence, played a role by providing unwanted visual references. We also wondered whether the aperture had provided some type of motion signal that could inform about the drift direction. For all these reasons, we repeated the experiment using a custom-built display, an array of 110 × 8 LEDs specifically selected to provide no persistence and no baseline luminance (Fig. 2A). We also made sure to rule out any possible motion signal by exposing each Vernier bar only for a brief interval (5 ms), the shortest detectable exposure allowed by our display.

**Fig. 2:**
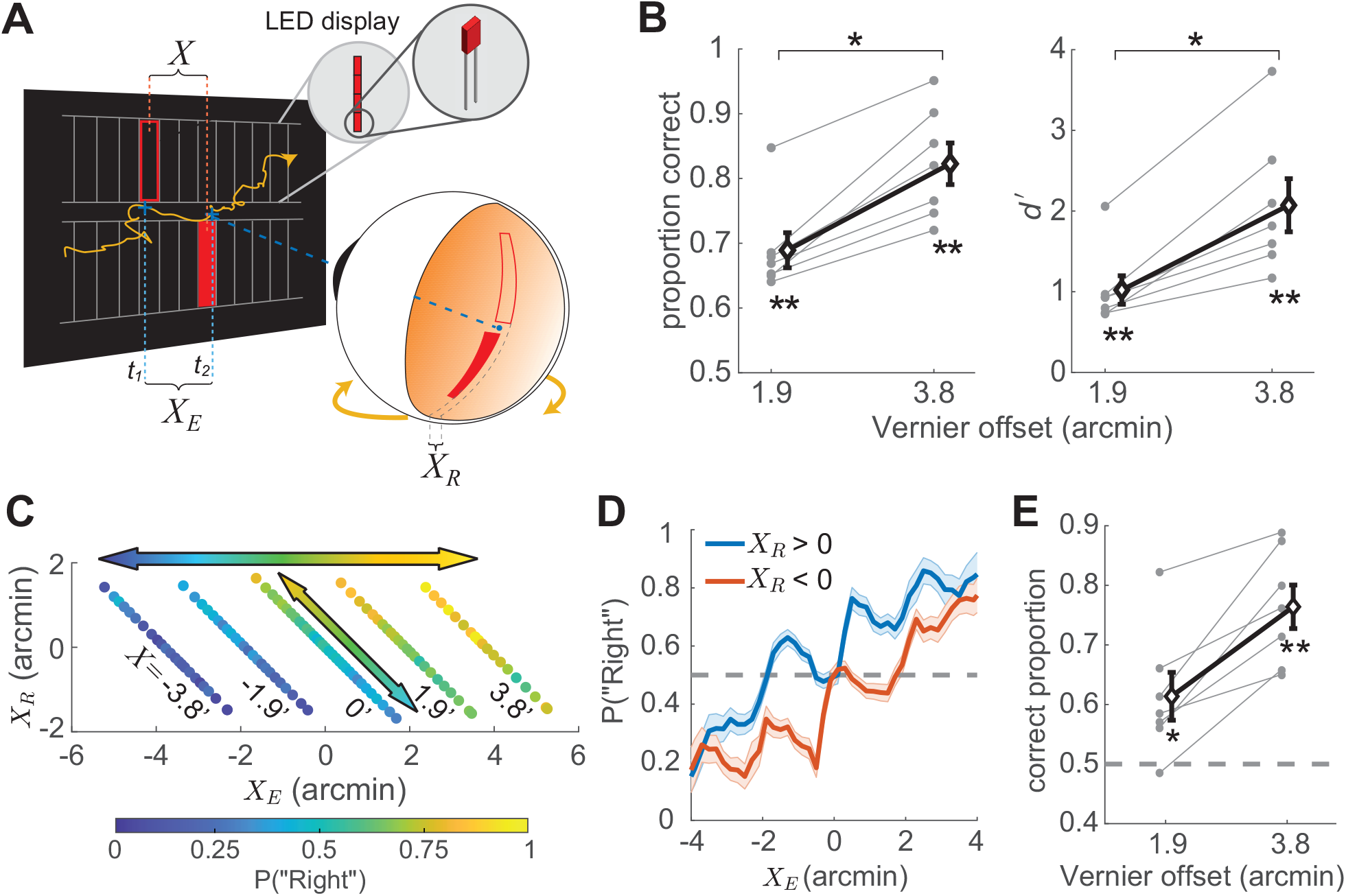
Controlling for visual cues. (**A**) A custom LED display developed specifically for this study. This display, an array of 110 × 8 LEDs, was designed to provide no persistence and no background luminance. Each LED covered a visual angle of 1.9′ on the horizontal meridian. (**B**) Performance measured with 5 ms exposures. Graphic conventions are as in Fig. 1H (*N* =7; ^*\**^*p <* 0.0017, paired two-tailed t-test; ^*\*\**^*p <* 0.0012, above chance, two-tailed t-test). **(C)** Probability of ‘Right’ responses as a function of both the eye displacement in a trial (*X*_*E*_) and the small misalignment on the retina caused by the display resolution (*X*_*R*_ *<* 1.9′; one LED, see panel *A*). Negative and positive *X*_*R*_ indicate that, on the retina, the bottom bar was shifted to the left or right, respectively. Each diagonal line represents a Vernier offset *X* on the display. **(D)** Marginal probability of ‘Right’ responses as a function of the eye displacement in a trial for both *X*_*R*_ *<* 0 and *X*_*R*_ *>* 0. The shaded regions represent one SEM. **(E)** Performance in the trials in which *X*_*E*_ and *X*_*R*_ possessed opposite signs. Subjects successfully completed the task even when *X*_*R*_ predicted the wrong response (^*\**^*p <* 0.03; ^*\*\**^*p <* 0.0004; above chance; two-tailed t-test).

Comparison between the data in Fig. 2B and Fig. 1H show that results were little affected by these changes in visual stimulation. The drift behavior changed little from the previous experiment and remained practically identical to that observed during fixation on a visible marker (Fig. S2). Critically, subjects continued to correctly report the stimulus even under these more stringent conditions: performance was already above chance at the smallest possible Vernier offset (in this case 1.9′, the width of one LED) and further improved as the distance between the two bars increased. These effects were clearly visible in the data from each observer, all of whom individually exhibited above chance performance at all Vernier offsets presented (*p <*0.011, bootstrap test).

These findings were very robust. As in the experiment of Fig. 1, performance was similar for leftward and rightward ocular drift (Fig. S1D-F), showing access to the specific drift trajectory performed in each trial, rather than knowledge of possible directional biases. Results were also not caused by possible inaccuracies in measuring eye movements. In this regard, it is important to notice that the experiments relied on the relative alignment of the two bars on the retina, not their absolute positions. That is, conclusions do not depend on the accuracy of gaze localization—a notoriously difficult operation—but on the capability to measure changes in gaze position, something that a properly tuned and calibrated DPI eye-tracker accomplishes with sub-arcminute resolution ^43^. Monte Carlo simulations show that eye movements would need to be over-estimated by an unrealistic amount, over 100%, to account for our findings (Supplementary Fig. S3). This degree of imprecision is not plausible with our recording apparatus.

Furthermore, analysis of residual errors in the alignment of retinal stimuli revealed that these cues cannot account for the experimental data. To be perfectly aligned on the retina, each bar needs to be rendered exactly at the current location of gaze. In practice, however, the precision of this operation is limited by the resolution of the display, as the stimulus can only be shown at the closest pixel/LED location, resulting in a small offset (*X*_*R*_ in Fig. 2A). This misalignment did exert a perceptual influence. For each Vernier offset *X* on the display (diagonal lines in Fig. 2C), perceptual reports exhibited a subtle but systematic influence from *X*_*R*_: the probability of reporting the bottom bar to the right was slightly larger when the misalignment was consistent with this interpretation (*X*_*R*_ *>* 0) than when it was in the opposite direction (*X*_*R*_ *<* 0; diagonal arrow in Fig. 2C). However, this cue could not possibly account for the general pattern of results obtained as *X* varied. Its influence was small relative to that exerted by the gaze displacement *X*_*E*_ (horizontal arrow in Fig. 2C), and overall, perceptual reports were driven by *X*_*E*_ irrespective of *X*_*R*_ (Fig. 2D). In fact, *X*_*R*_ was overall poorly correlated with subject responses (average correlation coefficient across observers: *ρ* = −0.016 ± 0.093), and subjects were able to successfully accomplish the task even in the trials in which the misalignment indicated the wrong response, the trials in which the *X*_*R*_ was in the opposite direction of the Vernier offset on the display (Fig. 2E). All these analyses further support the conclusion that humans incorporate fine oculomotor knowledge in the establishment of spatial representations.

The small stimulus offsets caused by the display resolution provide an opportunity to examine how the visual system integrates retinal and extraretinal signals at fixation. To gain insight into this process, we compared the perceptual reports recorded in the experiments to the responses of an ideal observer that inferred the most likely configuration of the stimulus from sensory measurements of both the eye displacement and the retinal misalignment. The ideal observer assumes uncertainty in sensory signals (modeled as additive Gaussian noise) and possesses only general knowledge about eye movements. Specifically, it assumes that ocular drift evolves as Brownian motion and, therefore, the variance of the probability of gaze displacement increases proportionally to time^44,45^. For each individual observer, the diffusion constant of this motion was directly estimated from their eye movements. In each trial, the model weighted the measured probability of eye displacement by its prior and estimated the most likely configuration of the stimulus (“Left” or “Right”) by comparing the overall probability (the integral of the 2D posterior probability distribution) on the two sides of the zero displacement line (diagonal cyan line in Fig. 3A).

**Fig. 3:**
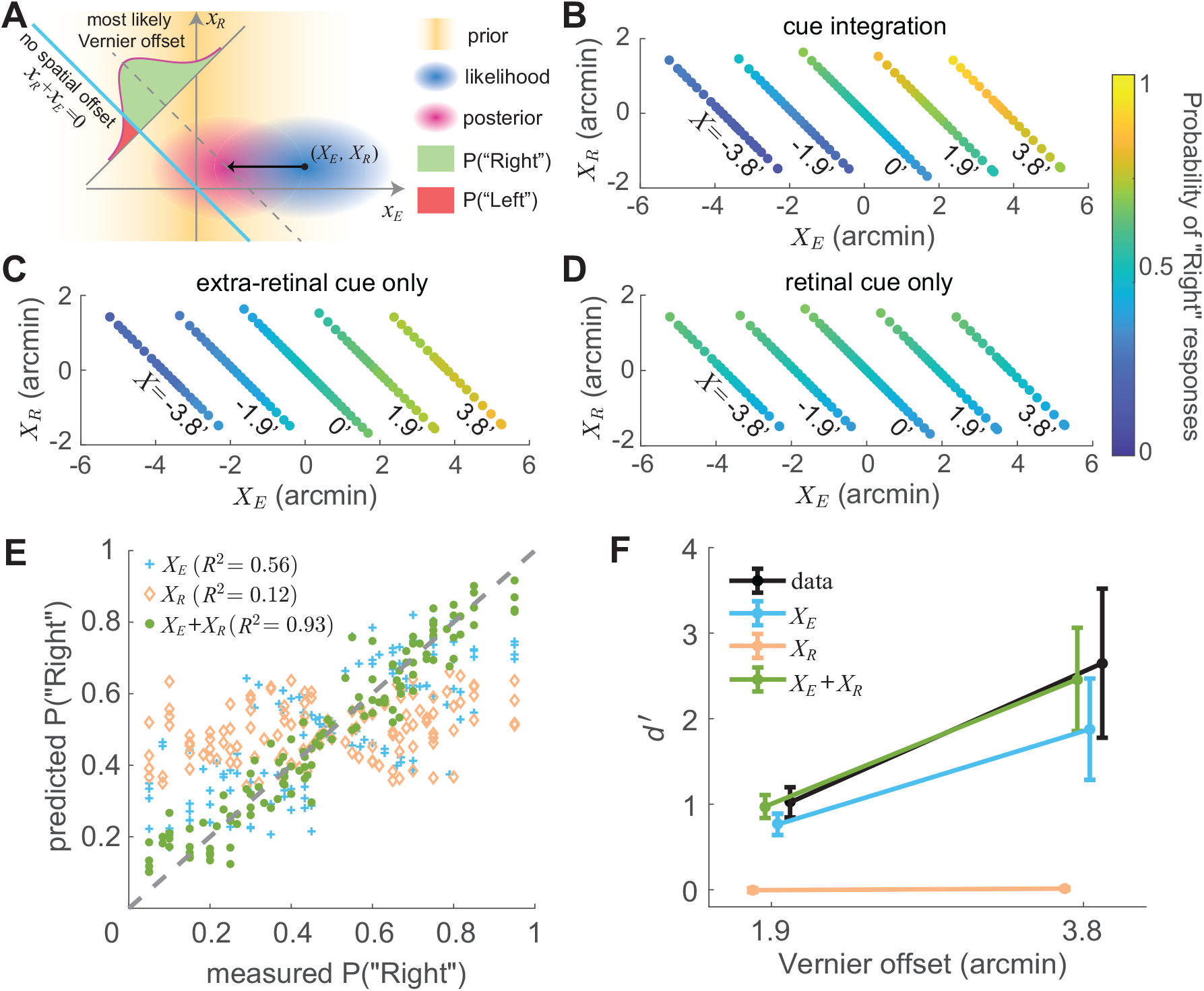
Integration of visual and motor cues. **(A)** An ideal observer model that combines retinal (*x*_*R*_) and extra-retinal (*x*_*E*_) signals. The model assumes sensory measurements to be corrupted by unbiased additive Gaussian noise (standard deviations *σ*_*R*_ and *σ*_*E*_) and applies a uniform prior to *x*_*R*_ and a zero-mean Gaussian prior to *x*_*E*_ (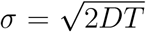, where *T* is the ISI), the latter based on the assumption that ocular drift resembles Brownian motion. In each trial, the likelihood of any given combination (*X*_*E*_, *X*_*R*_) is first converted into a joint posterior probability distribution and then integrated on the -45^*°*^ line *x*_*E*_ + *x*_*R*_ = *X* (dashed line) to evaluate the probability of any given Vernier offset *X*. The left/right perceptual report in a trial is determined by which side of the zero-offset line (cyan line) gives higher probability. **(B-D)** Response patterns are best predicted by the model when combining both cues (*B*); the model (*C*) that only uses extraretinal information does not account for the dependence on *X*_*R*_, and the model (*D*) that uses only retinal information fits poorly. Graphic conventions are as in Fig. 2*C*. **(E-F)** Comparison of experimental data and model predictions of responses and d′: (*E*) Probability of responding “Right” for various combinations of *X*_*R*_ and *X*_*E*_ (the same data points as in *B*-*D*). (*F*) Average performance measured as d′. Error bars represent ± one SEM across subjects. The cue integration model predicts both subject responses and overall performance with significantly greater accuracy than the single-cue models.

The ideal observer closely replicated the way subject’s responses varied as a function of *X*_*E*_ and *X*_*R*_ (*cf*. Fig. 3B and Fig. 2C). As in the empirical data, the overall pattern of response was primarily driven by the eye displacement, but a dependence on retinal misalignment was also visible for each gap of the Vernier on the display. Across all data points, the model accounted for 93% of the variance in subject’s responses (green dots in Fig. 3E) and accurately predicted the d′ observed in the experiments (green line Fig. 3F; individual data in Fig. S4A). Critically, both motor information about drift displacement and retinal information about bars alignment were necessary to replicate experimental data. Discarding the retinal signal led to a reduction in performance, but the model was still able to account for about 56% of the variance in perceptual reports. In contrast, performance dropped to chance level and the model could only account for 12% of the variance following elimination of extraretinal information (Fig. 3E,F; see also log-likelihood data in Fig. S4A). Thus, subjects performed very similarly to the predictions of a Bayesian combination of retinal and extraretinal sensory signals, with a predominant influence exerted by motor knowledge of eye movements.

The previous results indicate that motor knowledge of eye drifts during fixation is incorporated into spatial judgements. What are the mechanisms responsible for monitoring gaze position at this level of resolution? To gain insight into this process, we first examined the size and position of the temporal window over which gaze displacement best correlated with subject’s responses. As shown in Fig. 4*A*, the correlation peaked for a short window of approximately 100 ms that slightly preceded the bar presentations. This finding was highly consistent across individual observers, all of whom exhibited a similar timing, resulting in a statistically significant anticipation (−13 ms on average; Fig. 4*B*). Thus, during fixation, retinal signals appear to be combined with motor estimation of gaze position that slightly precedes retinal exposures, suggesting a predictive use of extraretinal signals.

**Fig. 4:**
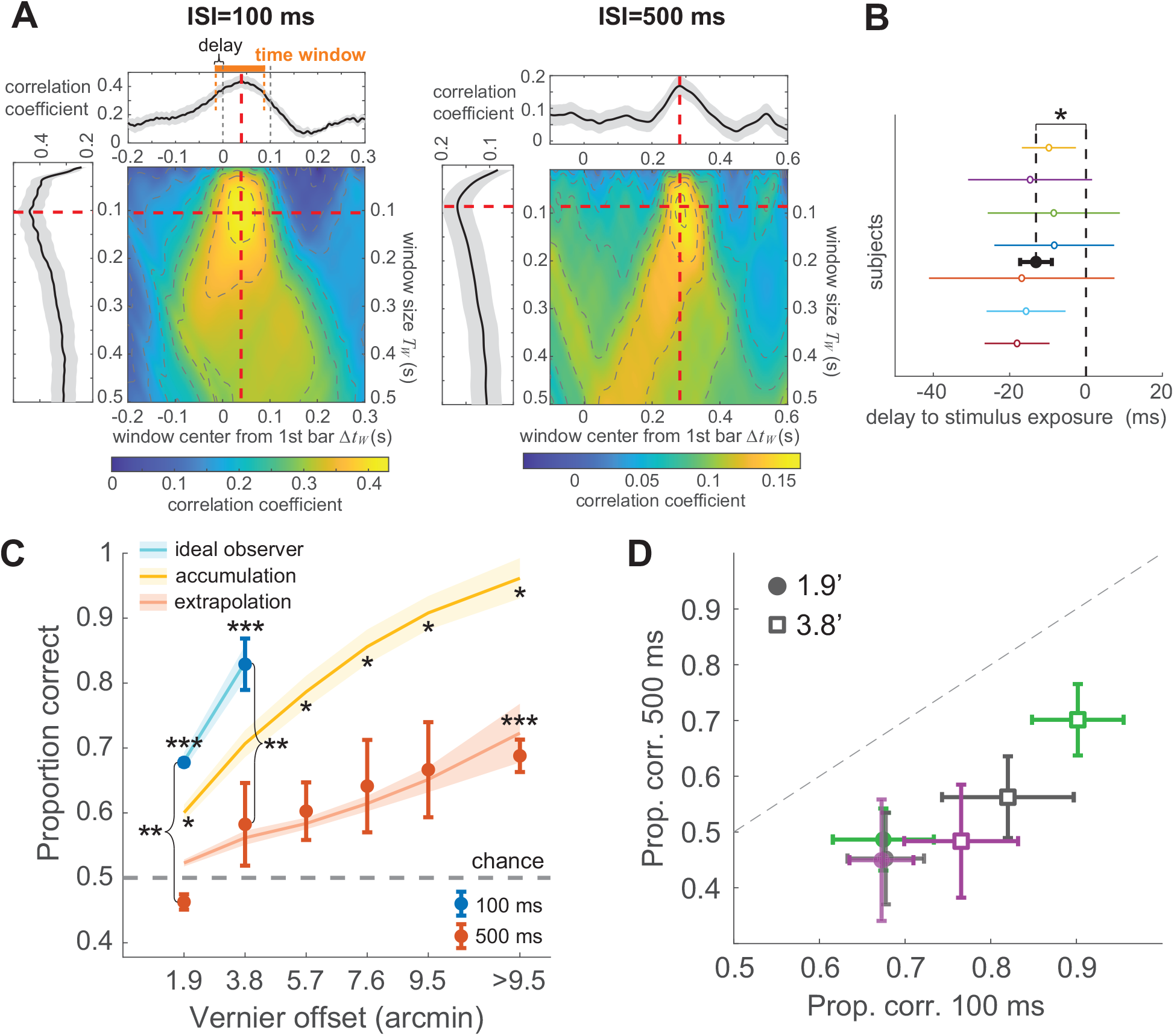
Characteristics of the extraretinal signal. **(A)** Correlation between perceptual reports and the horizontal gaze displacement 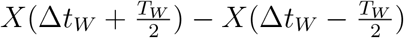 as a function of both window duration (*T*_*W*_ ; *y*-axis) and temporal lag between window center and bars exposure (Δ*t*_*W*_ ; *x*-axis). The highest correlation is achieved for a 100 ms window that slightly (13ms) precedes retinal stimulation. Side plots are sections at the optimal *T*_*W*_ and Δ*t*_*W*_ (red dashed lines). **(B)** The timing of maximum correlation for each individual subject. On average, subject responses are best correlated with a 100-ms window that anticipated the stimulus by 13 ms (filled black circle; ^*\**^*p <* 3.5 10^*−*4^, two-tailed t-test). Error bars represent one SD. **(C)** Comparison of performance with two different ISIs: 100 and 500 ms. Data were collected using the custom LED display with 5 ms flashes (N=3). Performance was lower in the 500 ms condition (^*\*\**^*p <* 0.005, paired one-tailed t-test) and improved marginally with increasing Vernier offset (^*\*\*\**^above chance; *p <* 0.009, one-tailed t-test). Empirical data are consistent with the prediction from the ideal observer model with *σ*_*E*_ adjusted to increase proportionally to time (red curve) and lower than predicted by increasing 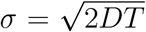 (yellow curve; ^*\**^*p <* 0.037, one-tailed paired t-test). Note that these fits have no free parameters: all parameters were obtained from those estimated over the 100 ms ISI in Figure 3. Error bars and shaded regions represent ± one SEM. **(D)** For each observer (different colors) performance at both 1.9′ and 3.8′ gaps was always lower in the 500 ms condition (*p <* 0.027, one-tailed bootstrap). Error bars represent ± one SEM.

Given that the duration of the optimal window in Fig. 4 was similar to the interval between bar exposures (100 ms), we wondered whether this window indicates continuous oculomotor monitoring throughout the ISI or represents a fixed internal temporal scale over which drift is estimated. Both possibilities can be mediated by a mechanism of integration of noisy velocity signals, a process similar to the one believed to occur for smooth pursuit ^46–48^. However, these two hypotheses lead to distinct predictions as the ISI is further increased. If drift displacement is integrated across the entire interval between bar exposures, we would expect the uncertainty in the extraretinal measurement of displacement (*i*.*e*., its standard deviation *σE*) to increase no faster than 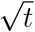, as accumulation of temporally uncorrelated noise progressively disrupts the position estimate. In contrast, if gaze displacement is estimated from the movement measured over a shorter interval of the ISI, we would expect *σ*_*E*_ to increase proportionally to *t*, as the consequence of a temporal extrapolation process.

To address this question, we repeated the experiment while increasing the ISI between the bar exposures by a factor of five, to 500 ms. Except for this longer interval, the paradigm was otherwise identical to that of Fig. 2, with 5 ms exposures delivered by our custom LED display. Increasing the ISI profoundly affected performance. Proportion of correct responses at the smallest Vernier offsets dropped drastically and were at now chance level with a 1.9′ gap. Furthermore, even at the much larger Vernier offsets resulting from the longer ISI, performance remained considerably lower than the levels measured in the 100 ms ISI condition (Fig. 4C). Results were highly consistent across subjects, all of whom exhibited substantial and significant reductions in performance in the 500 ms condition (Fig. 4D).

These data closely matched the predictions of our ideal observer model under the “extrapolation” hypothesis, *i*.*e*., when the uncertainty in the extraretinal measurement for the 500-ms ISI was assumed to be five times larger than for the 100-ms ISI (*σ*_*E*_(500) = 5 *σ*_*E*_(100); red curve in Fig. 4C). In contrast, model predictions fell far from the data under the “integration” hypothesis, *i*.*e*., when extraretinal uncertainty was increased proportionally to the *E*squared root of time (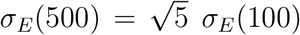; yellow curve in Fig. 4C). In this case, the model significantly overestimated performance in all observers (Fig. S5). In keeping with these data, the duration of the temporal window over which gaze displacement best correlated with perceptual reports did not increase, but remained similar to that observed for the shorter ISI (Fig. 4A). These findings are not compatible with continuous monitoring of eye position throughout the ISI. They suggest that extraretinal monitoring of ocular drift is conducted over a short window and the displacement over a longer interval is estimated via a process of extrapolation.

## Discussion

The eyes drift incessantly in the intervals between saccades, even when attending to a single point, raising fundamental questions on how the visual system avoids perceptual blurring, resolves fine detail, and establishes stable high-acuity spatial representations. Existing theories assume these processes to rely exclusively on the spatiotemporal luminance input to the retina^6,12,49,50^. Contrary to this idea, our results show that humans have access to high-resolution motor knowledge about eye movements and integrate this information with signals from the retina to estimate fine spatial relations. These findings challenge the standard view of passive processing of a retinal image during fixation and indicate that the computations responsible for representing visual space are intrinsically sensorimotor.

To unveil extraretinal contributions, our study relied on methods for gaze-contingent display control, the updating of the stimulus according to the observer’s eye movements. This approach enables both precise control of visual input signals and manipulation of visuomotor contingencies. Specifically, in our experiments, we tailored the visual input to create a stimulus configuration on the display that conveyed no spatial information on the retina. Extensive care was taken to eliminate all informative spatial cues in the retinal input. Precautions included: (a) presenting stimuli in complete darkness, and (b) preventing dark adaptation to avoid sensitivity to possible spurious light sources; (c) using a specifically designed custom display that was not affected by visual persistence nor baseline luminance; and (d) using very brief exposures to eliminate motion signals. Furthermore, (e) use of stationary stimuli and tight head immobilization, together with (f) selective consideration of trials in which the eye exclusively moved because of ocular drift, ensured that our findings were not contaminated by corollary discharges associated to saccades, microsaccades, or other types of smooth eye movements.

Our data show that even under these stringent conditions, humans retain sufficient knowledge of their oculomotor activity to reconstruct the direction in which gaze drifts over a short interval. This knowledge is specific: it is continually collected over a short temporal interval of approximately 100 ms, enabling detection of gaze displacements with arcminute resolution. This window of observation systematically anticipates the visual input by approximately 10 ms, likely yielding an even larger anticipation once neural delays are taken into account. Furthermore, this oculomotor signal is evaluated in the light of general knowledge of eye drift statistics, so that spatial judgements closely follow the predictions of an ideal observer that assigns uncertainty to the estimated spatial representation (the addition of retinal and extraretinal information) based on the reliability of the extraretinal signal.

It is important to emphasize that our results cannot plausibly be explained by inaccurate positioning of stimuli on the retina nor inaccuracies in eye-tracking. Our analyses confirm that our apparatus is highly precise, as we can reliably measure the perceptual consequence of the stimulus misalignment caused from display resolution, the tiny mismatch between the measured gaze position and the actual position of the stimulus on the display. While this retinal cue exerts a clear influence at every presented Vernier offset, it cannot account for perceptual reports, as performance in the task varied primarily with the measured gaze displacement. Modeling of eye-tracking errors showed that gaze displacement would need to be over-estimated by an unrealistic amount to yield retinal cues that could account for our results. Such level of imprecision is not plausible with the DPI eye-tracker used in these experiments. This system was periodically tested during the course of data collection to ensure that it accurately measured one-arcminute rotations of an artificial eye. Furthermore, measurements errors are expected to go in the opposite direction in an analog DPI, as the inertia of the moving components would tend to underestimate—rather than amplify—very small eye movements.

It is also worth pointing out that stimulus duration was too short to provide useful motion signals. The 5 ms exposure used in the experiments of Figs. 2-4 is about one order of magnitude shorter than that necessary to perceive motion in the speed range of ocular drift^51,52^. The median displacement of the stimulus on the retina during such brief interval was only ∼ 14 arcseconds, well below the thresholds reported in the literature for similar tasks^53^, and results did not change when selecting only the trials with minimal displacement during exposure. In keeping with these observations, the instantaneous velocities measured around the times of bar exposures were only weakly correlated with perceptual reports (Fig. 4A), providing further evidence that retinal image motion played no role in our experiments.

The finding that eye drift is monitored appears to contradict widespread assumptions in the field. An obvious conflict is with the notion that drift is not controlled—the popular idea that this motion results from noise at the neural and/or muscular level^32,54^. Although less known, however, it has long been proposed that the smooth fixational motion of the eye actually represents a form of slow control, a sort of pursuit of a stationary target aimed at maintaining ideal visual conditions^37,38,55^. This view has received strong support in the recent literature. It is now known that during natural fixation, when the head is free to move normally, ocular drift partially compensates for the physiological instability of the head, severely constraining retinal image motion^39,56^. Furthermore, changes in the characteristics of ocular drift have been observed in high-acuity tasks, as when looking at a 20/20 line of an eye-chart or when judging the expression of a distant face ^40,57^. These changes appear to be functional, as they increase the power of the luminance modulations impinging onto retinal receptors, an effect consistent with theories arguing for temporal representations of fine spatial details^9,10^. The present study goes beyond this previous body of work by showing that the signals involved in exerting control at this scale also contribute extraretinal information that is integrated in spatial representations.

Our conclusions also contrast with those reached by previous studies with similar paradigms. Classical experiments with asynchronously displayed Verniers concluded that drift is not monitored because performance declines with increasing delays between exposures^34,58^. Fig. 4 replicates this effect, but our data show that other factors (*e*.*g*., memory decays and/or the window over which drift is monitored) must be responsible for the measured decrement in performance. More recently, support to the notion that fixational drift is not monitored has come from systematic localization errors observed with stimuli briefly displayed in complete darkness. When reporting the position of a previously displayed reference by selecting between two probes, one at the same reference’s location on the display (spatiotopic probe) and one at its same position on the retina (retinotopic probe), subjects systematically select the retinotopic one^35^. These errors are, in fact, predicted by our ideal observer model, but attributed to the specific perceptual choice presented to the observer rather than lack of extraretinal knowledge of eye drift (see Fig. S6). Thus, the present study suggests alternative explanations for the previous reports in the literature.

Our findings lead to a critical question: why are eye movements monitored at such high level of resolution? There are several complementary ways in which an extraretinal drift signal could contribute to visual processing. A possibility is by facilitating visual stability during fixation, *i*.*e*., by helping disentangling the visual motion signals resulting from external objects from those generated by eye movements. Studies on how the visual system discards motion signals resulting from egomotion have primarily focused on larger eye movements, saccades and smooth pursuits^28,29,31,59,60^. These studies have emphasized the interaction between retinal and extraretinal signals, both efference copies of motor commands ^61,62^ and proprioceptive information from extraocular muscles ^63,64^. The eye drift that occurs during fixation is commonly assumed to be too small for extraretinal compensation, and the resulting visual motion signals are believed to be perceptually canceled solely on the basis of the retinal input^36^. This idea, however, is at odds with the motion perceived during exposure to retinally-stabilized objects, stimuli that move with the eye to remain immobile on the retina^35,65^. Furthermore, it has been observed that motion perception is biased to the direction of eye movements, so that stimuli that move opposite to ocular drift on the retina tend to appear stable even if their motion is amplified^65,66^. Such bias requires knowledge of drift direction, information that could be provided by the extraretinal signal uncovered in our experiments.

In this regard, it should be observed that the jitter after-effects that follow adaptation to dynamic noise patterns do not speak against contributions from extraretinal signals. Local adaptation to a selected portion of the visual field yields a perceived jittery motion of a stationary pattern in the complementary, unadapted region. This phenomenon seems to exclude the possibility that eye motion is solely estimated from an extraretinal signal, as subtraction of retinal and eye velocities would leave residual motion in the adapted region rather than the unadapted one. However, the jitter after-effect is compatible with estimation of drift motion via a combination of both retinal and extraretinal signals. For example, the visual system could estimate drift motion as the lowest instantaneous velocity signal on the retina that is also congruent with drift direction, a view that would explain not only jitter after-effects, but also the perceived motion with stabilized images and the directional anisotropy in motion perception. Note that this approach differs from a purely retinal cancellation mechanism for also requiring directional consistency with extraretinal measurements.

Our findings suggest another way in which extraretinal drift information could contribute, which is by directly participating in the establishment of high-acuity visual representations. In our experiments, observers were able to infer geometrical arrangements purely based on extraretinal information. Until now, spatial information during fixation has been assumed to be extracted solely from the visual signals impinging onto the retina ^6,12,49,50^. While several methods have been proposed for registering afferent visual information into spatial maps as the eye drifts, all these methods exclusively rely on the retinal input. However, this process presumably depends on the richness of visual stimulation and requires temporal accumulation of evidence, difficulties that an extraretinal drift signal could alleviate. Thus, motor knowledge of ocular drift may be particularly valuable when visual stimulation is sparse and following saccades, when new visual content is introduced on the retina. Interestingly, an extraretinal contribution makes this process similar to the coordinate transformation underlying the establishment of head-centered spatial representations during larger eye movements^67–71^, emphasizing a general computational strategy and supporting a similarity between fixational drift and pursuit movements^38^. Further work is needed to assess the origins of the extraretinal signal unveiled by our experiments and its specific role in representing space.

## Methods

### Subjects

A total of 13 subjects (5 males and 8 females; age range: 20-35), all naïve about the purpose of the study, participated in the experiments. All subjects were emmetropic, with at least 20/20 visual acuity in the right eye as measured by a Snellen eye chart, and were compensated for their participation. Informed consent was obtained from all participants following the procedures approved by Institutional Review Boards at Boston University and the University of Rochester.

### Stimuli

Stimuli consisted of standard Verniers, with two vertical bars separated by a horizontal gap (the Vernier offset; Fig. 1A). The two bars were never simultaneously visible: they were exposed at different times at the current location of the line of sight on the display, so that the offset was determined by the gaze displacement that occurred in between the two exposures. In this way, the bars always appeared vertically aligned on the retina, whereas the gap on the display varied across trials based on the eye movements performed by the observer. Each bar was 28′ long and 1.4′ wide in the experiment of Fig. 1 (Experiment 1), and 19′ × 1.9′ in the experiments of Figs. 2 and 4 (Experiments 2 and 3). These dimensions were the outcome of adjusting the distance of the display so that each bar could be as thin as possible (one pixel wide in Experiment 1 and one LED wide in Experiments 2-3), while at the same time retaining clear visibility when briefly exposed at maximum intensity. Stimuli were examined in total darkness, carefully removing all light sources that could serve as potential spatial references and all visual cues that could provide information about the Vernier configuration.

### Apparatus

Stimuli were rendered by means of EyeRIS, a hardware/software system for real-time gaze-contingent display that enables precise synchronization between eye movement data and the refresh of the display^72^. They were viewed monocularly with the right eye, while the left eye was patched. A dental imprint bite-bar and a headrest minimized head movements and maintained the observer at a fixed distance from the monitor.

Different displays were used in Experiment 1 and in Experiments 2-3. In Experiment 1, stimuli were rendered on a fast-phosphor CRT monitor (Iiyama HM204DT) at a resolution of 800 × 600 pixels and 200 Hz refresh rate. This monitor has fast phosphors with decay time shorter than 2 ms. A completely dark background and tuning of the monitor at minimum settings ensured that the edges of the display were never visible.

To further control for possible influences from phosphor persistence, residual background luminance, and retinal image motion, in Experiments 2-3 stimuli were displayed on a custom LED display specifically developed for this study (Fig. 2A). LEDs are not affected by lingering activity like phosphors and have zero baseline illumination when not active. The custom display consisted of 880 LEDs, 800 rectangular elements arranged into two rows of 4 LED, and a 3 × 3 array of circular LED used for eye-tracking calibration. Each Vernier bar was given by the simultaneous activation of a column of 4 LED in either the top or the bottom row. This display also offered lower latency relative to a CRT (3 ms vs. 7.5 ms, on average) and more precise timing, since each LED could be controlled independently without having to wait for the rasterization of a frame to be completed, as in a CRT. LED activation triggered a digital signal that was sampled synchronously with oculomotor data, so that the timing of stimulus presentation could be reconstructed offline with high precision.

To measure eye movements with the precision necessary to align stimuli on the retina, we used a dual Purkinje Image (DPI) eye-tracker (Fourward Technology), an analog system with high spatiotemporal resolution and minimal delay. This specific eye-tracker has been customized over the course of two decades to refine its dynamics and minimize sources of noise. It resolves movements smaller than 1′ as tested with an artificial eye controlled by a galvanometer. Analog eye movements data were first low-pass filtered at 500 Hz, then sampled at 1 kHz, and recorded for off-line analysis.

### Experimental procedures

Data were collected in multiple experimental sessions, each lasting approximately 1 hour. Each session consisted of several blocks of trials, with each block containing approximately 50 trials. Every block started with preparatory procedures to ensure optimal eye-tracking. These steps included positioning the subject in the apparatus, tuning the eye-tracker, and performing calibration procedures to accurately localize gaze. Frequent breaks between blocks allowed the subject to rest. Lights were turned on during these breaks to prevent dark adaptation and minimize visibility of the edges of the CRT as well as the influence of any possible residual light.

Subjects were told that the two bars of a Vernier would be presented sequentially in random order and were asked to report whether the bottom bar was to the left/right of to the top bar by pressing a corresponding button on a joypad. Each trial started with the subject fixating on a 5′ red dot at the center of the display for 1 s. The fixation marker was then turned off, and after a uniformly-distributed random delay of 1-2 s, the first Vernier bar was exposed either above or below the current gaze position (equal probability across trials). The second bar then followed with a fixed delay (the inter-stimulus interval, ISI) at the current gaze location. In this way, the two Vernier bars were aligned on the retina and separated on the display by the gaze shift that occurred during the ISI (both horizontal and vertical displacements in Experiment 1; only horizontal displacement in Experiments 2 and 3). The ISI was 100 ms in Experiments 1 and 2 and 500 ms in Experiment 3.

Slightly different procedures were adopted in Experiments 1 and 2-3. In Experiment 1, the image was continually updated on the CRT display to replicate the visual consequences of viewing stimuli through a thin slit aperture that moved with gaze (*i*.*e*., a retinally-stabilized aperture; Fig. 1C,D). This implied that the stimulus exposure varied across trials, as each bar remained visible as long as it was aligned with the aperture. One bar was displayed in the top half of the aperture, and one in the bottom half. In Experiments 2 and 3, to eliminate possible motion signals, each Vernier bar was only displayed for 5 ms, the shortest exposure at the maximum intensity afforded by our LED display. In every trial, two columns of LED were activated, one in the top and one in the bottom row of the display. Columns were selected as the ones closest to the horizontal gaze position measured at the time of exposure. Except for these points, the paradigm was otherwise identical in the two experiments.

### Data analysis

#### Oculomotor data

Periods of blinks and poor tracking were automatically detected by the DPI eye-tracker. Only trials with optimal, uninterrupted eye-tracking and no blinks were selected for data analysis. Recorded oculomotor traces were first automatically segmented into separate periods of drift and saccades based on speed threshold of 3^*°*^*/s* and validated by human experts. Segmentation based on eye speed is very accurate with the high-quality data provided by the DPI during head immobilization. In this study, we specifically focus on ocular drift. All trials that contained other types of eye movements besides ocular drift, like saccades and microsaccades, were excluded from data analysis.

#### Evaluation of performance

At every Vernier offset, performance was quantified by means of both proportion correct and d′. For each individual observer, we used bootstrap to evaluate statistical significance across conditions and differences from chance levels (Figs. S4 and S5). The data reported in Figs. 1-4 are averages across observers and corresponding statistics.

In Fig. 2, performance is also examined as a function of both the horizontal eye displacement (*X*_*E*_) and the estimated misalignment of the two bars on the retina (*X*_*R*_ *<* 2′). Ideally the two Vernier bars need to be perfectly aligned. However, each bar could only be displayed at the pixel/LED closest to the estimated gaze position, so that the Vernier offset *X* on the display was not *X*_*E*_, but equal to *X*_*E*_ + *X*_*R*_. We assessed the joint influence of *X*_*E*_ and *X*_*R*_ by binning trials according to their values to uniformly sample the space and examined how perceptual reports varied across bins. In the space (*X*_*E*_, *X*_*R*_), a Vernier offset *X* on the display corresponds to a -45^*°*^ tilted line, as the same *X* could be reached with various cue combinations (*X*_*E*_ + *X*_*R*_ = *X*). The 5 lines in Fig. 2*C* corresponds to the 5 Vernier offsets reached in the experiment (0′, ±1.9′, ±3.8′). The data in Fig. 2*C* represent averages obtained by pooling data across subjects, so that each bin contained on average 60 trials.

In Fig. 4*A, B*, the correlation between gaze displacement and perceptual reports was examined as a function of both lag Δ*t*_*W*_ and duration *T*_*W*_ of the temporal window of observation. To this end, we first converted subject’s responses into a binary format (−1 and 1) and then computed the Pearson correlation coefficient with the horizontal displacement in the interval 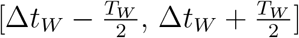.

#### Ideal observer model

To gain insight into the mechanisms by which extraretinal estimation of ocular drift contributes to representing space, we compared the perceptual reports measured in the experiments to those of an ideal observer that adds noisy sensory measurements of spatial cues on the retina and eye movements (*X*_*R*_ and *X*_*E*_) to establish head-centered representations. The ideal observer assumes ocular drift to resemble Brownian motion with a specific diffusion rate. This assumption is incorporated in the joint prior distribution *p*(*X*_*R*_, *X*_*E*_), which is uniform along *x*_*R*_ and follows a Gaussian distribution with zero mean and standard deviation 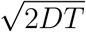 along *x*_E_, where *D* is the diffusion coefficient of the individual’s drift process and *T* the ISI. In a Brownian process the variance evolves proportionally to time. For each subject, we estimated *D* from the recorded eye traces via linear regression of the variance of the gaze displacement over the considered ISI. Sensory measurements of *X*_*E*_ and *X*_*R*_ were assumed to be corrupted by independent additive white noise processes with Gaussian distributions: *p*(*x*_*R*_ | *X*_*R*_) = *N*(*X*_*R*_, *σ*_*R*_) and *p*(*x*_*E*_ | *X*_*E*_) = *N*(*X*_*E*_, *σ*_*E*_). In every trial, the ideal observer estimates the joint posterior probability of the retinal and extraretinal displacement:

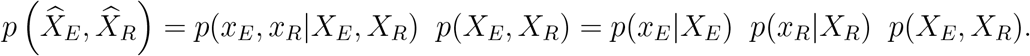

Thus, 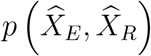 is a two-dimensional Gaussian with mean and covariance given by:

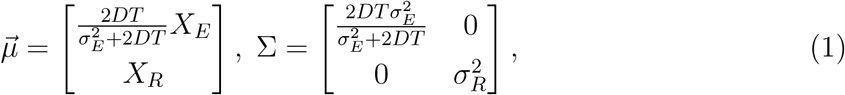

The probability of any given Vernier offset 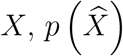, can then be estimated by integrating the joint posterior probability 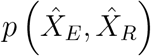 along the line *X*_*E*_ + *X*_*R*_ = *X* (Fig. 3A):

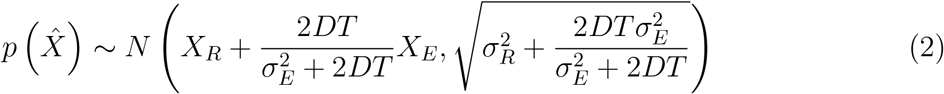

The probabilities of reporting the bottom bar of the Vernier to the left or to the right of the top bar are then given by 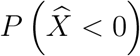 and 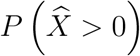, respectively.

The two free parameters of the model, *σ*_*E*_ and *σ*_*R*_, determine the uncertainty of sensory measurements. The larger is *σ*_*E*_, the weaker is its perceptual influence, with no trial-specific extraretinal knowledge of ocular drift in the limit case of *σ*_*E*_ = ∞. These parameters were estimated individually for each subject to maximize the log likelihood 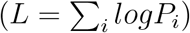 of the model replicating the subject’s perceptual reports across all trials:

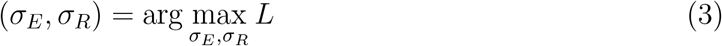

where *P*_*i*_ represents the probability that the model responds in the same way as the observer in trial *i*: 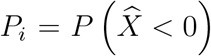 if the subject responded “Left” and 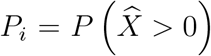 if he/she responded “Right”.

#### Evaluation of model performance

We evaluated the model in several ways. The data in Fig. 3*F* compare the overall performance measured in the experiments to that predicted by the model. Predictions were first obtained for each individual observer (Fig. S4*A*) and then averaged across subjects in Fig. 3. The log-likelihood *L* by which the model accounts for subject’s perceptual responses is reported in Fig. S4*A*. We also examined the model’s capability to reproduce the pattern of perceptual responses as a function of the measured retinal and extraretinal cues. Fig. 3*B* compares the output of the model to perceptual reports for each of the groups of trials of Fig. 2*C*. The overall accuracy of the model is summarized by the coefficient of determination *R*^2^ in Fig. 3*E*.

Furthermore, in Fig. 3 we compared both performance and perceptual reports to an ideal observer that operates on just one of the two cues, either *X*_*E*_ or *X*_*R*_. In this case, parameters were optimized with the model reduced to estimating the Vernier offset from the marginal posterior probability along the considered axis:

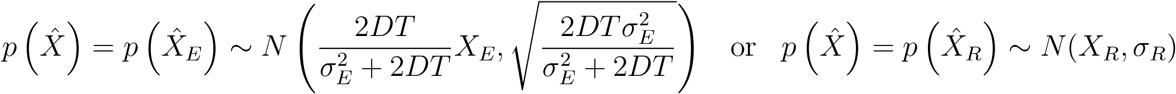

where parameters were obtained via the same maximum likelihood procedure used for the full model.

#### Dynamics of drift estimation

Distinct predictions emerge if gaze displacement is estimated over the entire interval between bar exposures or by extrapolating measurements obtained over a shorter interval. In the former case, the error in estimating gaze displacement will progressively accumulate because of the noise in the measurement. Specifically, the standard deviation of the estimate will grow as 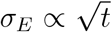 under the assumption of temporally uncorrelated sensory noise. In contrast, if drift is estimated over an interval shorter than the ISI, we would expect the displacement error to grow proportionally to time as a consequence of extrapolation: *σ*_*E*_ ∝ *t*.

In Fig. 4*C*, we tested which of these two alternative hypotheses best fit the data when the ISI, *T*, was increased from 100 ms to 500 ms. In the 500 ms condition, the standard deviation of the prior was correspondingly increased by a factor of 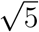 to reflect the five-fold increment in the interval between bar exposures, as dictated by the assumption that ocular drift resembles Brownian motion. The uncertainty in the retinal signal (*σ*_*R*_) remained the same as in the 100 ms condition. The uncertainty in the extraretinal cue (*σ*_*E*_) was either enlarged by a factor a 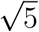 or 5 as suggested by the two hypotheses. Individual subjects data and model predictions are reported in Fig. S5.

## Acknowledgements

This work was supported by the National Institutes of Health grants EY18363 and EY07977. We thank Claudia Cherici and David Richters for their help in preliminary experiments and Janis Intoy and Martina Poletti for helpful comments and discussions.

## Supplementary Information

**Fig. S1:**
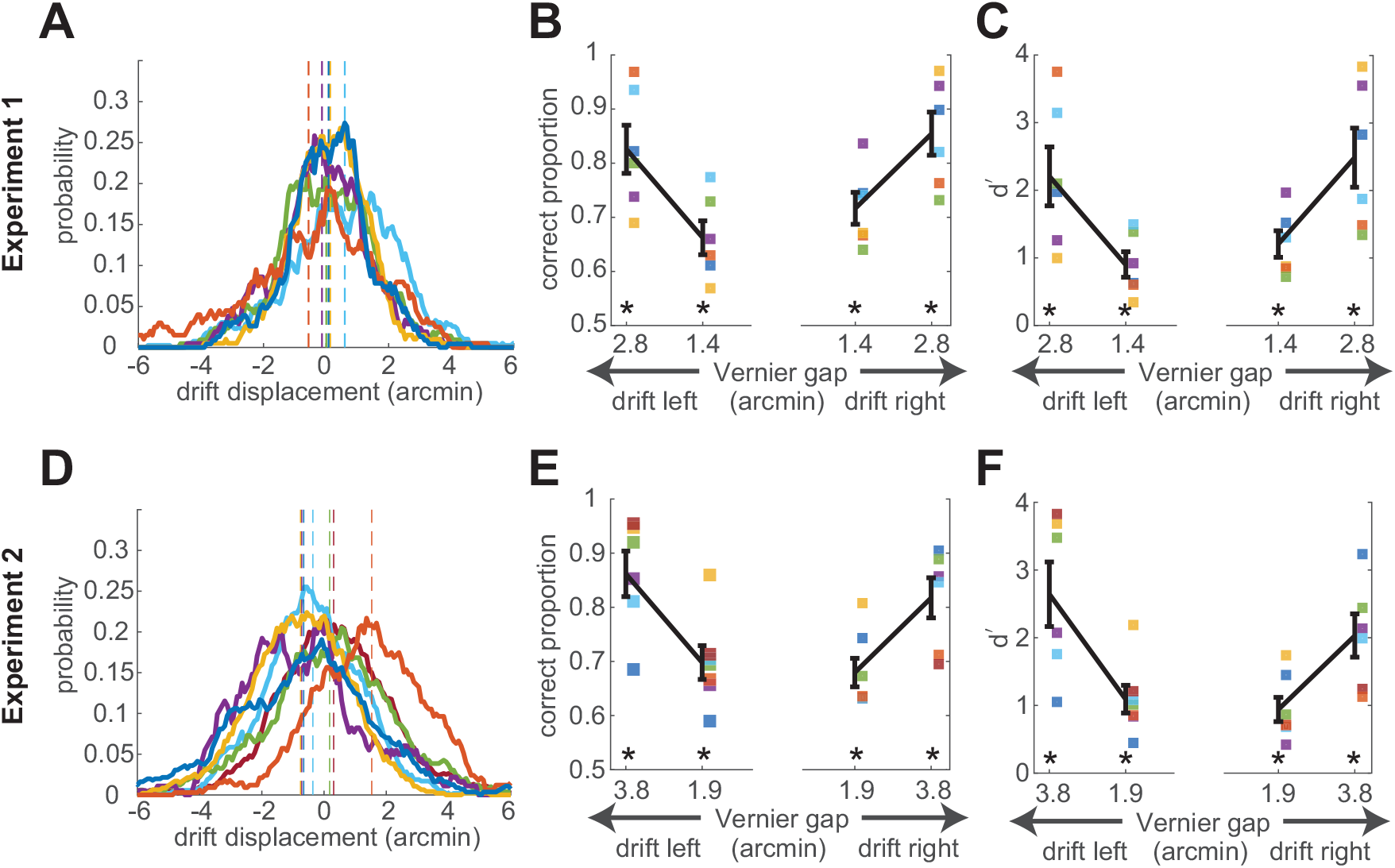
Performance as a function of ocular drift direction. **(A)** Distributions of horizontal eye displacement in the 100 ms inter-stimulus interval of Experiment 1. Data from individual subjects are shown in separate curves. The vertical dashed lines mark the means of the distributions. Note that for all observers means are close to zero, *i*.*e*., drift displacements were unbiased. **(B-C)** Performance in Experiment 1 measured as both proportion correct (*B*) and *d*′ (*C*) for displacements in both directions. Black lines represent averages across subjects ± one SEM. Squares are data from individual subjects (^*\**^*p <* 0.0008 in *B* and *p <* 0.0052 in *C*, two-tailed t-tests). Results with drifts in both directions were similar. **(D-F)** Similar analyses for the data from Experiment 2. Graphic conventions are identical to the panels above.

**Fig. S2:**
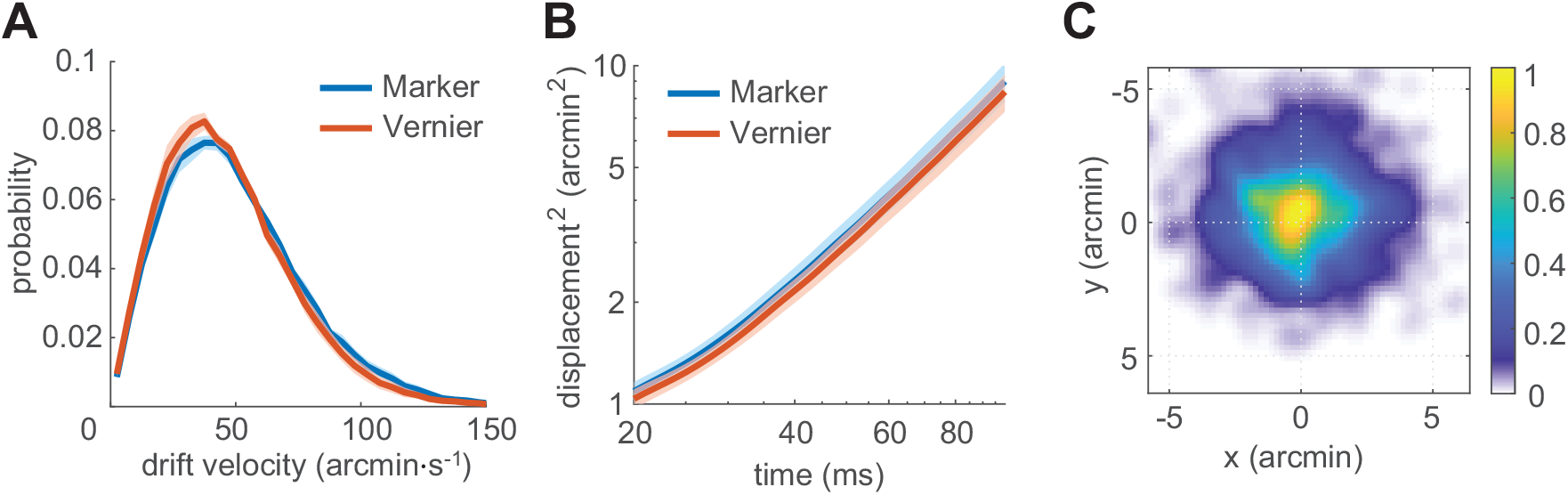
Ocular drift characteristics in Experiment 2. (**A**) Average distribution of eye speed. (**B**) Squared displacement as a function of time. (**C**) 2D probability of overall drift displacement. Data represent averages across individuals and refer to the 100 ms ISI interval. Shaded regions represent ± one SEM. For comparison, the same measurements obtained while maintaining fixation on a 5′ dot (marker) are also shown in *A* and *B*.

**Fig. S3:**
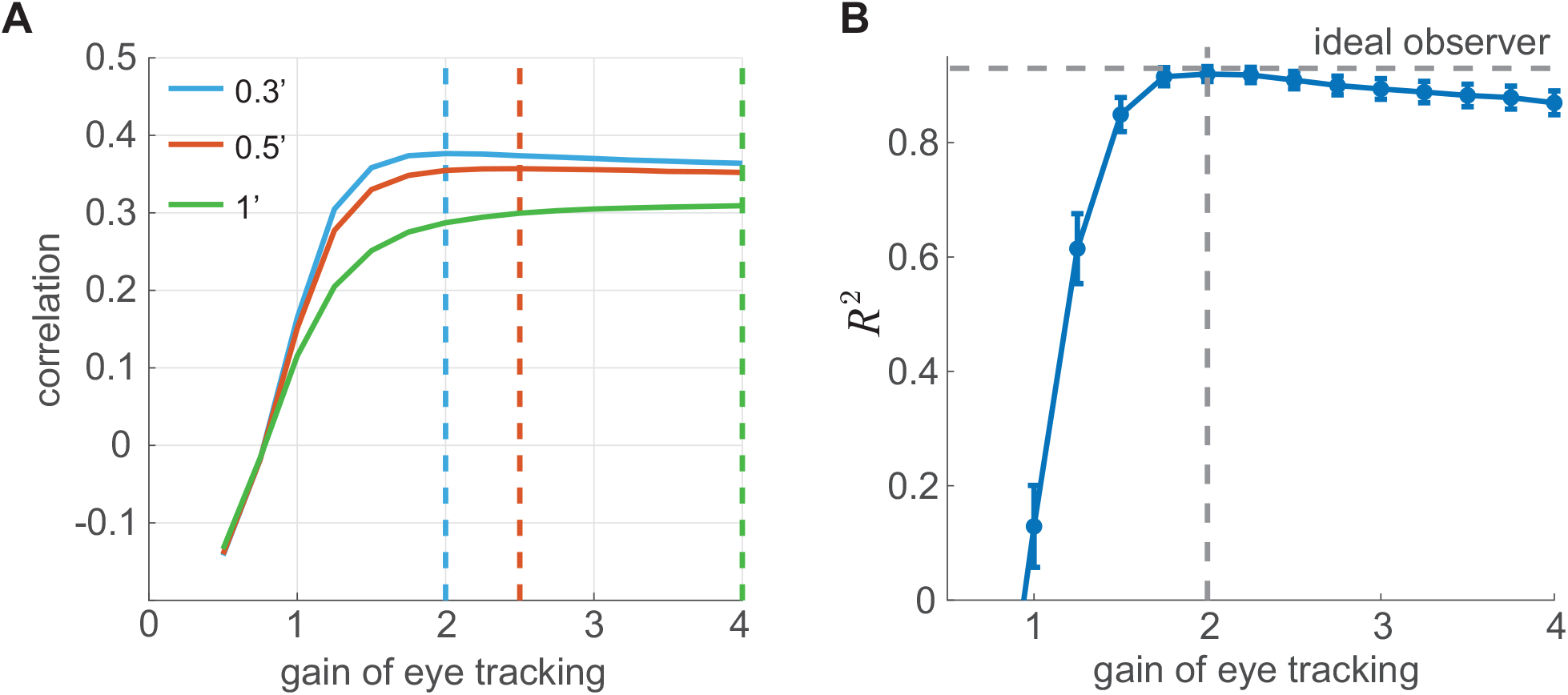
Consequences of over-estimating eye movements. Results of Monte Carlo simulations that modeled the eye-tracker output as 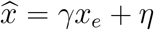, where *x*_*e*_ is the horizontal gaze displacement; *γ* represents the eye-tracker gain; and *η* = *N* (0, *σ*) is a Gaussian noise term with zero mean and standard deviation *σ*_*η*_. (**A**) Correlation between subject’s responses and the resulting retinal misalignment 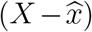 as a function of *γ*. The three curves represent results with different *σ*_*η*_. The lower boundary for *σ*_*η*_, as measured with a stationary artificial eye is 0.3. Note that the correlation never exceeds 0.4. Vertical dashed lines show the gains for which the curves reach their maximum. **(B)** Maximum variance in perceptual reports that could be explained by an ideal observer only using this retinal cue. For each *γ*, the perceptual uncertainty in the retinal measurement was estimated to maximize the *R*^2^ as in Eq. 3. To account for subject’s responses, the eye-tracker would need to overestimate the gaze displacement by approximately a factor of 2 (vertical line), which is unrealistic. The dashed horizontal line marks the variance accounted by the ideal observer in Figure 3, which assumes measurements of eye drifts to be veridical (*γ* = 1).

**Fig. S4:**
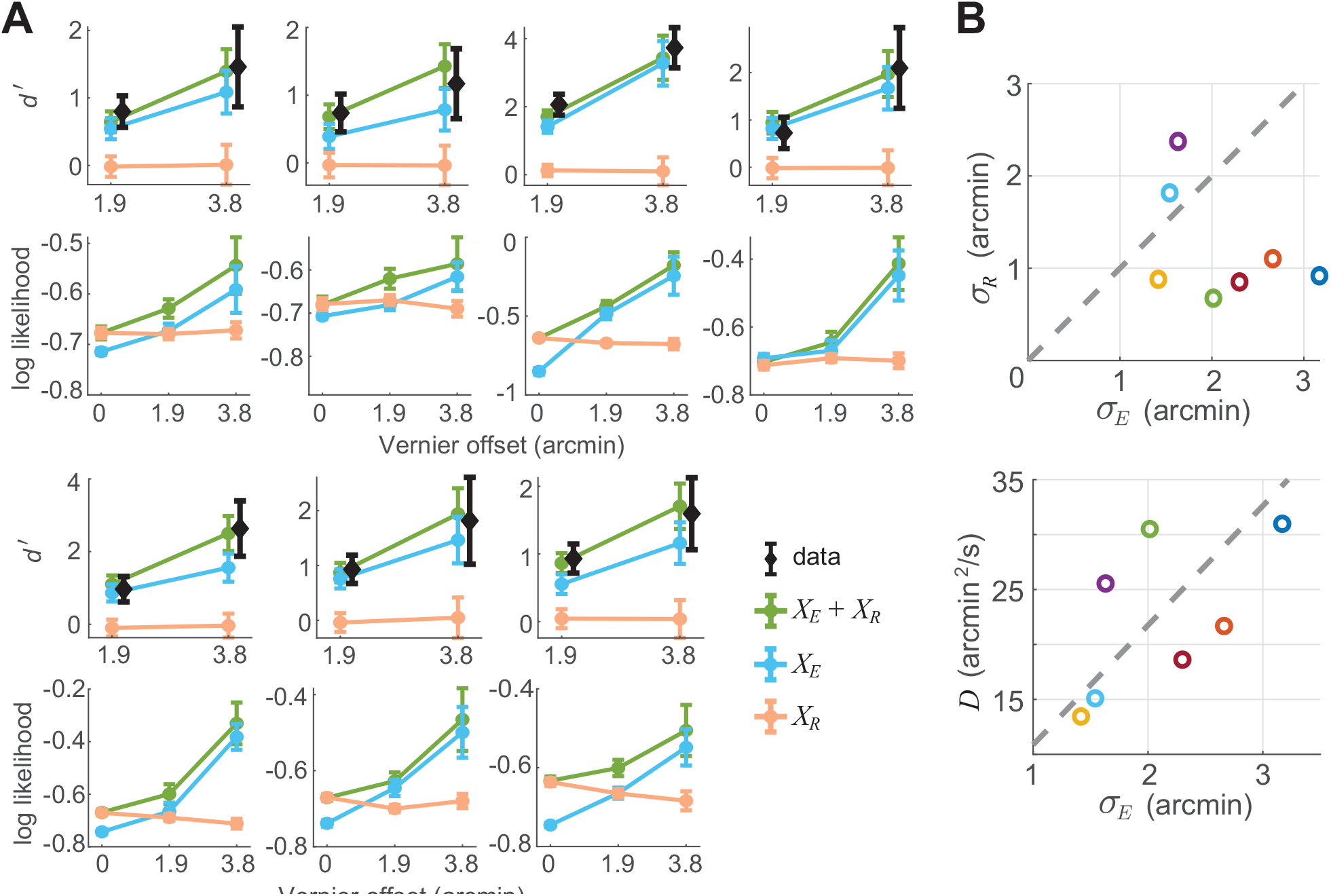
Model parameters and predictions for individual subjects. **(A)** For each subject (N=7) in Experiment 2, 3 models are compared, including the full Bayesian model (*X*_*R*_ + *X*_*E*_) and the reduced model with single cue (*X*_*R*_ and *X*_*E*_). Their performance was evaluated via the comparison between the predicted d’ and empirical one (top row) and the mean log likelihood of all trials given the model (bottom row. *L* in the Methods; see Eq. 3). Errorbars are ± one SEM derived from bootstraps. **(B)** Model parameters fitted to empirical data in Experiment 2. Each data point corresponds to one subject: The s.d. of retinal noise *σ*_*R*_ and uncertainty in extraretinal displacement estimation *σ*_*E*_ in the top panel; The diffusion coefficient of ocular drift in the bottom panel.

**Fig. S5:**
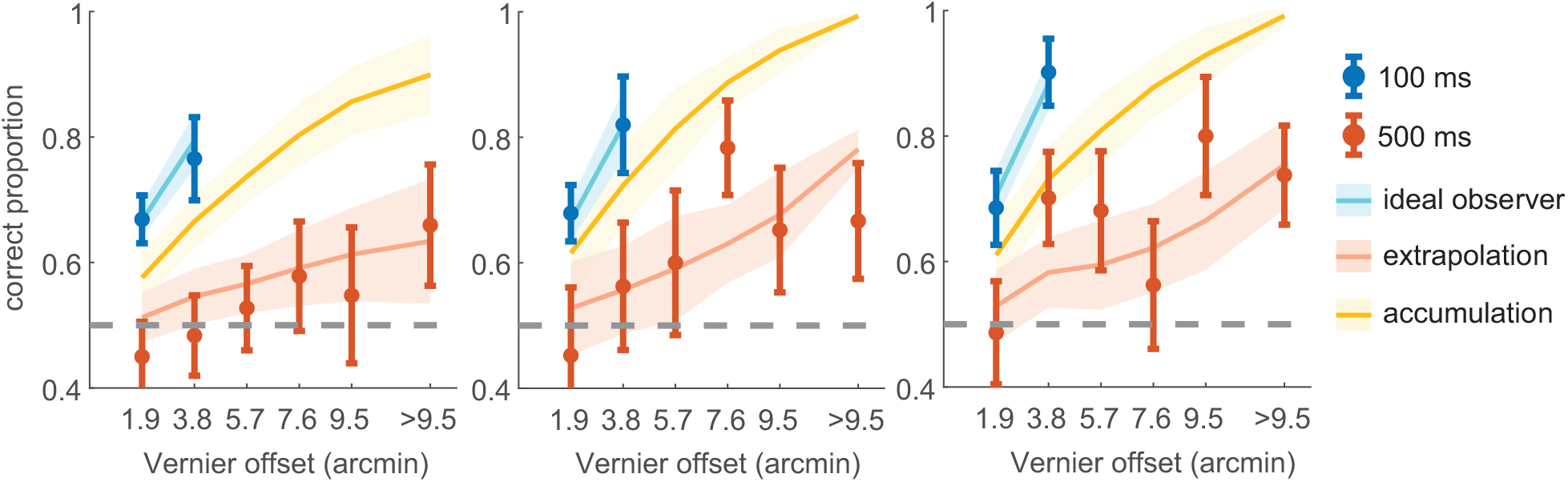
Model predictions for individual subjects in Experiment 3. The graphical convention is the same as Fig. 4C. Error bars of the empirical data and the shaded region of model predictions are ± one SEM derived from bootstraps.

**Fig. S6:**
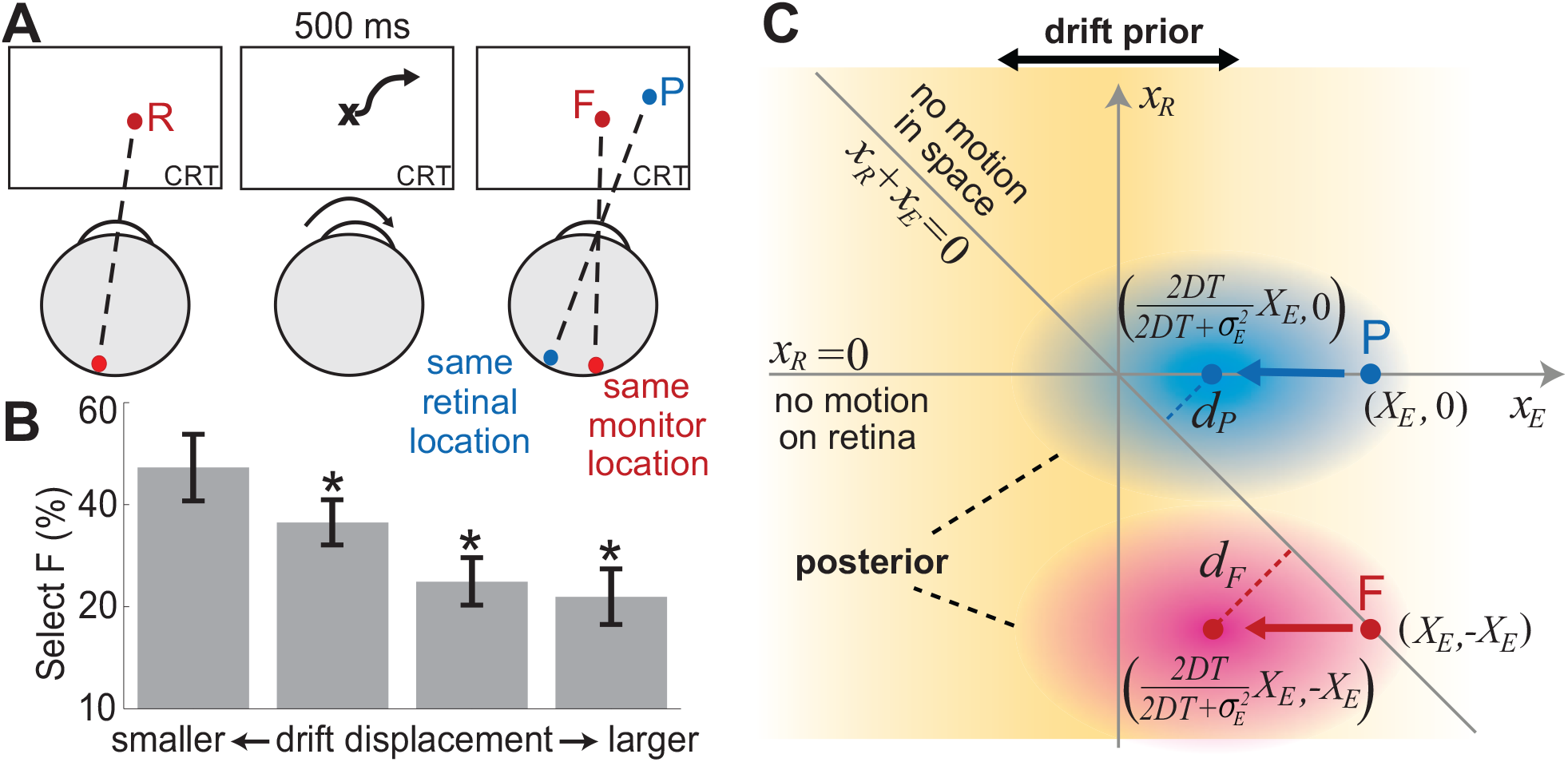
Predicted errors in spatial localization. Our ideal observer model accounts for seemingly contradictory previous findings. (**A**) In a 2AFC task, subjects report the position of a previously displayed reference (*R*) by selecting between two probes, one at the same reference’s location on the display (*F*) and one at its same position on the retina (*P*). (**B**) The more the eye drifts in complete darkness, the less likely subjects are to correctly select *F* (data from [35], reprinted with permission). (**C**) The model predicts this paradoxical behavior as a consequence of the specific choice presented to the observer. The oculomotor prior weights identically both probes, causing both posterior distributions to shift towards smaller estimated displacements. The posterior distribution of the retinotopic probe *P* will be closer to the no-motion line (the line *X*_*R*_ + *X*_*E*_ = 0) if the motor uncertainty in measuring the displacement, *σ*_*E*_, is larger than the variance of the prior (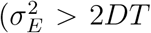, where *D* is the drift diffusion constant and *T* the ISI). The data in Fig. 4 confirm that this will occur for sufficiently long ISI. Under these conditions, the model predicts that the retinotopic probe *P* will have higher probability to be mistaken for the reference than the spatiotopic probe *F*, despite having access to an extraretinal drift signal.

